# Embryo Lethality Assay for Evaluating Virulence of Isolates from Bacterial Chondronecrosis with Osteomyelitis in Broilers

**DOI:** 10.1101/2021.03.12.435217

**Authors:** Nnamdi S. Ekesi, Amer Hasan, Alia Parveen, Abdulkarim Shwani, Douglas D. Rhoads

## Abstract

We used an embryo lethality assay (ELA) to assess virulence for different isolates from cases of bacterial chondronecrosis with osteomyelitis (BCO) in broilers. ELA has been used to measure virulence and lethal dosage of *Enterococcus faecalis* and *Enterococcus cecorum*. We hypothesized that ELA could substitute for more laborious and costly assessments of BCO isolate pathogenicity using live birds. We evaluated two different levels of bacteria injected into eggs from layer and commercial broiler embryos. Significant findings include a) *Escherichia coli* from neighboring farms operated by the same integrator had very different embryo lethality, b) isolate *Staphylococcus agnetis* 908 had low virulence in ELA, even though this isolate can induce more than 50% BCO lameness, c) *Enterococcus cecorum* 1415 also had low pathogenicity; even though it was recovered from severe bilateral tibial dyschondroplasia, d) human and chicken isolates of *S. aureus* had significant pathogenicity, e) virulence for some isolates was highly variable possibly corresponding with quality of the embryos/fertile eggs used, and f) ELA pathogenicity was much lower for our BCO isolates than previous reports which may reflect maternal environment. Overall, ELA virulence and BCO virulence are not always concordant indicating that that ELA may not be an effective measure for assessing virulence with respect to BCO.

**Importance:** Lameness is among the most significant animal welfare issues in the poultry industry. Bacterial infections are a major cause of lameness and different bacterial species have been obtained from lame broilers. Reliable lab-based assays are required to assess relative virulence of bacteria obtained from lame broilers. Embryo Lethality Assays have been used to compare virulence. Our results suggest that this assay may not be an effective measure of virulence related to lameness.

## Introduction

Bacterial chondronecrosis with osteomyelitis (BCO) is the leading cause of lameness in rapidly growing broilers (1–4). Lameness in broilers is significant as an animal welfare issue, and as a financial cost, in the poultry industry (2). Our research group isolated and characterized an isolate of *Staphylococcus agnetis*, designated 908, from lame broilers on our research farm (1). *S. agenetis* 908 is closely related to isolates from sub-clinical mastitis in dairy cows (5). *S. agnetis* 908 can induce greater than 50% BCO lameness by 56 days of age when administered in a single dose in drinking water at 10^4^ to 10^5^ CFU/mL on day 20 (1, 3, 6, 7). Our current model for lameness etiology is that environmental stress can lead to increased leakage (translocation) of bacteria across the gut and pulmonary epithelia into the blood system (1–3, 8–10). Particular species that are able to survive in the blood stream may colonize the growth plate, a vulnerable niche in the blood system of the rapidly growing leg bones of fast-growing broilers (2, 3, 11). Distinct bacterial species have been isolated from lame birds including *Staphylococcus aureus*, *Enterococcus cecorum*, and *Escherichia coli* (12–32). However, there are few comparisons of different BCO-associated species, or isolates, for pathogenicity (18, 33). In this study, we investigated the pathogenicity of BCO isolates of *S. agnetis*, *Staphylococcus chromogenes*, *E. coli*, *E. cecorum*, and *S. aureus* using an embryo lethality assay (ELA). The isolates were obtained from BCO lesions on our research farm or commercial broiler farms in Arkansas. ELA has been used to correlate the expression frequency of nine virulence-associated *E. coli* genes with embryo mortality (34). Borst et al. (33) used this technique to compare the virulence of *E. cecorum* isolated from broiler spinal lesions (kinky back) to non-pathogenic *E. cecorum* strains isolated from ceca of unaffected birds. Blanco et al. (35) used ELA to determine the virulence and the lethal dose of *Enterococcus faecalis*.

## Methods

### Microbiology

Isolation and handling of the isolates has been described (1, 36). Media included: CHROMagar Orientation (CO; DRG International, Springfield Township, NJ), tryptic soy broth (Difco brand, Becton, Dickinson and Company, Franklin Lakes, NJ); and Luria broth (LB; per liter 10 g tryptone, 5 g yeast extract, 5 g NaCl).

### Embryo Lethality Assay

Fertilized eggs were obtained from leghorns (LCL) and Cobb700 commercial broilers (BCL) on the University of Arkansas research farm. The eggs were washed with warm soapy water containing about 1% household bleach. Eggs were incubated (NatureForm^™^ Hatchery Systems, Jacksonville, FL, USA) at 37.5°C, a relative humidity of 56%, on autorotate. On day 12, stationary-phase bacterial cultures were diluted 1:200 in 1x phosphate buffer saline (1xPBS; 150mM NaCl, 10 mM potassium phosphate pH 7.2). CFU concentration was computed from Absorbance at 650 nm using pre-calibrated standard curves for each isolate, and then diluted in 1xPBS to the required concentration. Eggs were candled, and fertile eggs were injected using a tuberculin syringe and 20G needle (Becton, Dickinson and Company) with 100μL of the appropriate bacterial suspension, or vehicle control, into the allantois cavity as described (33). The opening was sealed with transparent box tape. Inoculated embryos were scored for mortality every day for 4 days after bacteria administration (33, 35).

### Statistical Analysis

The results of the ELA were analyzed with either Pearson’s Chi-squared (χ2) or Fisher’s Exact (FE) analysis using SAS and R software (SAS Institute. 2011; RStudio Team. 2016). Significant differences were accepted at P < 0.05.

## Results

### Embryo Lethality Assay with BCO isolates

To establish a suitable assay for comparing different isolates, we first injected *E. coli* 1413 a dilution series (10^3^ – 10^8^ CFUs) in sterile 1xPBS to estimate the lethal dosage for Leghorn Chicken Line (LCL) embryos (Figure 1). Injections of 1413 above 10^5^ CFUs had embryo lethality of 80 to 100%. We therefore assessed different BCO isolates at 10^5^ and 10^6^ CFUs (Table 1). We included *S. agnetis* 908 recovered from a femoral BCO lesion on our research facility as this isolate can induce lameness ≥50% by day 56 when administered in drinking water for two days to 20-day old broilers (1, 3, 6, 7). Surprisingly, 908 injections of even 10^6^ CFUs resulted in only 14% embryo lethality, a level not statistically different from 1xPBS control treatment (Figure 2A). For the methicillin-sensitive human *S. aureus* isolate 1302, originally retrieved from a wound (Table 1), injections of 10^5^ or 10^6^ CFUs resulted in 80% embryo lethality (Figure 2B). *Staphylococcus chromogenes* 1401 was recovered from an infected T4 vertebra of a chicken with “kinky back” (Table 1). Injections of 10^6^ CFUs resulted in only 7% embryo death, less than the 1xPBS control for that experiment (Figure 2C). *E. coli* 1409 was recovered from a tibial head necrosis lesion (Table 1). Injection of 10^5^ to 10^6^ CFUs resulted in no lethality through day 4 (Figure 2D). *E. coli* 1413 was isolated from the blood of a lame bird with bilateral BCO of the tibiae and femorae, where *E. coli* was also recovered from multiple lesions (Table 1). As before, injections of 10^5^ or 10^6^ CFUs into LCL resulted in approximately 80% embryo lethality (Figure 2E). *E. cecorum* 1415 was isolated from a tibial head abscess in a case of bilateral tibial dyschondroplasia (Table 1). ELA results for 10^5^ CFU showed slightly more lethality than 10^6^ CFUs but neither was statistically different from the PBS control (Figure 2F). We used two isolates (1510 & 1514) of *S. aureus* obtained from BCO lesions from two different birds in a commercial broiler house lameness outbreak (Table 1; Ekesi 2020). Draft genome assemblies for these isolates were highly related. The isolates showed different ELA results with 1510 lethality of 60% for 10^6^ CFU, while 1514 produced 47% lethality. Only the 1510 results were statistically different from the 1xPBS control (Figure 2G & 2I). *E. coli* 1512 and 1527 were recovered from the left and right femoral lesions of the same bird (Table 1; Ekesi, 2020). Draft genome assemblies for both 1512 and 1527 were determined to be virtually identical. ELA results for 1512 yielded 87% lethality for 10^6^ CFUs and 52% lethality for 10^5^, but only the 10^6^ results were statistically different from the 1xPBS control (Figure 2H). Therefore, the only isolates that showed significant lethality using LCL embryos were the human isolate *S. aureus* 1302, and chicken isolates *E. coli* 1413, *S. aureus* 1510, and *E.coli* 1512.

**Figure 1.**
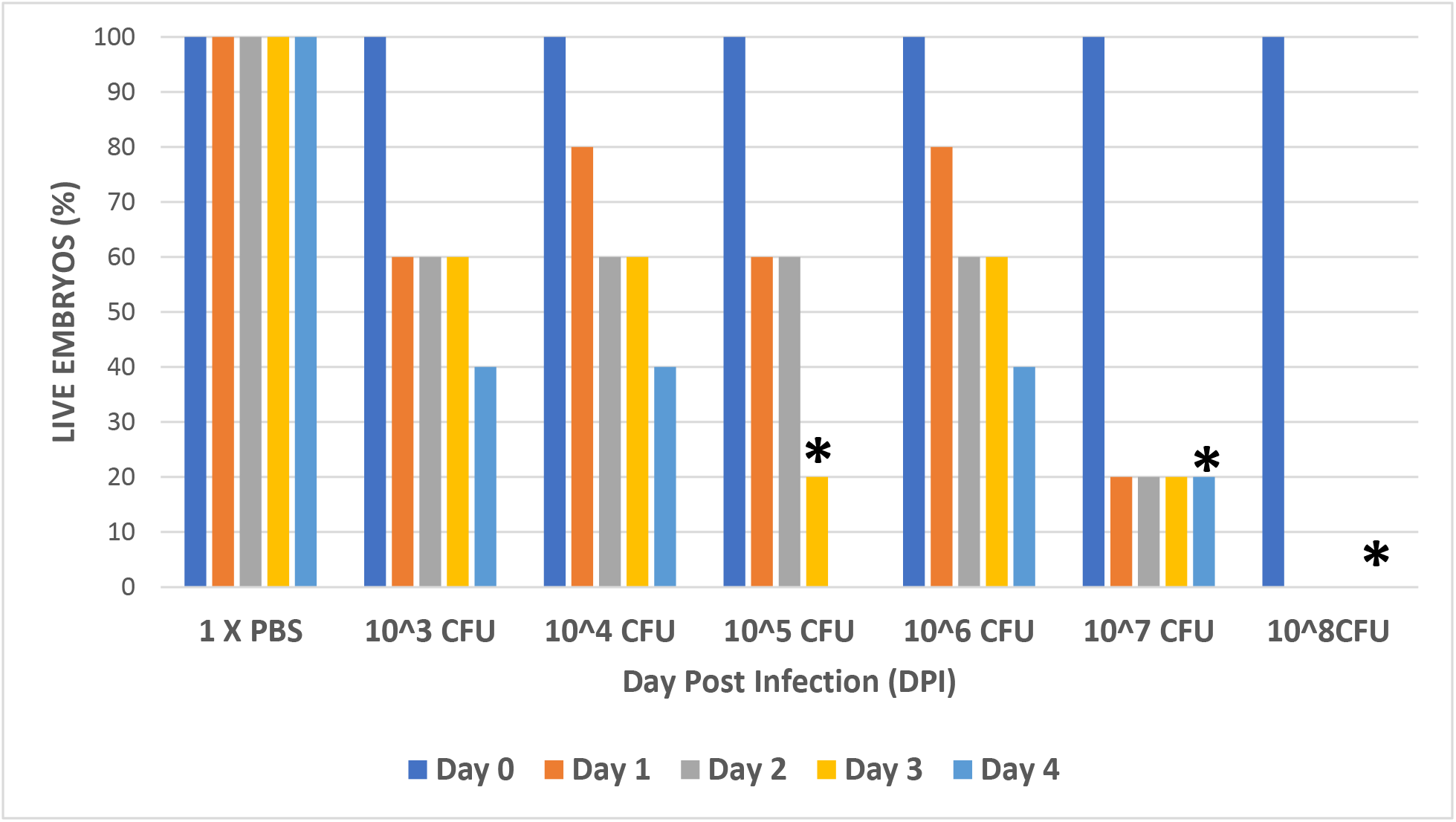
Layer chicken line embryo lethality for day post injection (DPI) with 1xPBS, different numbers of colony forming units (CFU) of *E. coli* 1413. The percentage (n=5) of live embryos (Y-axis) for different doses is graphed for four days post-injection (X-axis). Isolate source is described in Table 1. Asterisk (*) indicates where treatment was significantly different from 1xPBS (P <0.05) on day 4.

**Figure 2.**
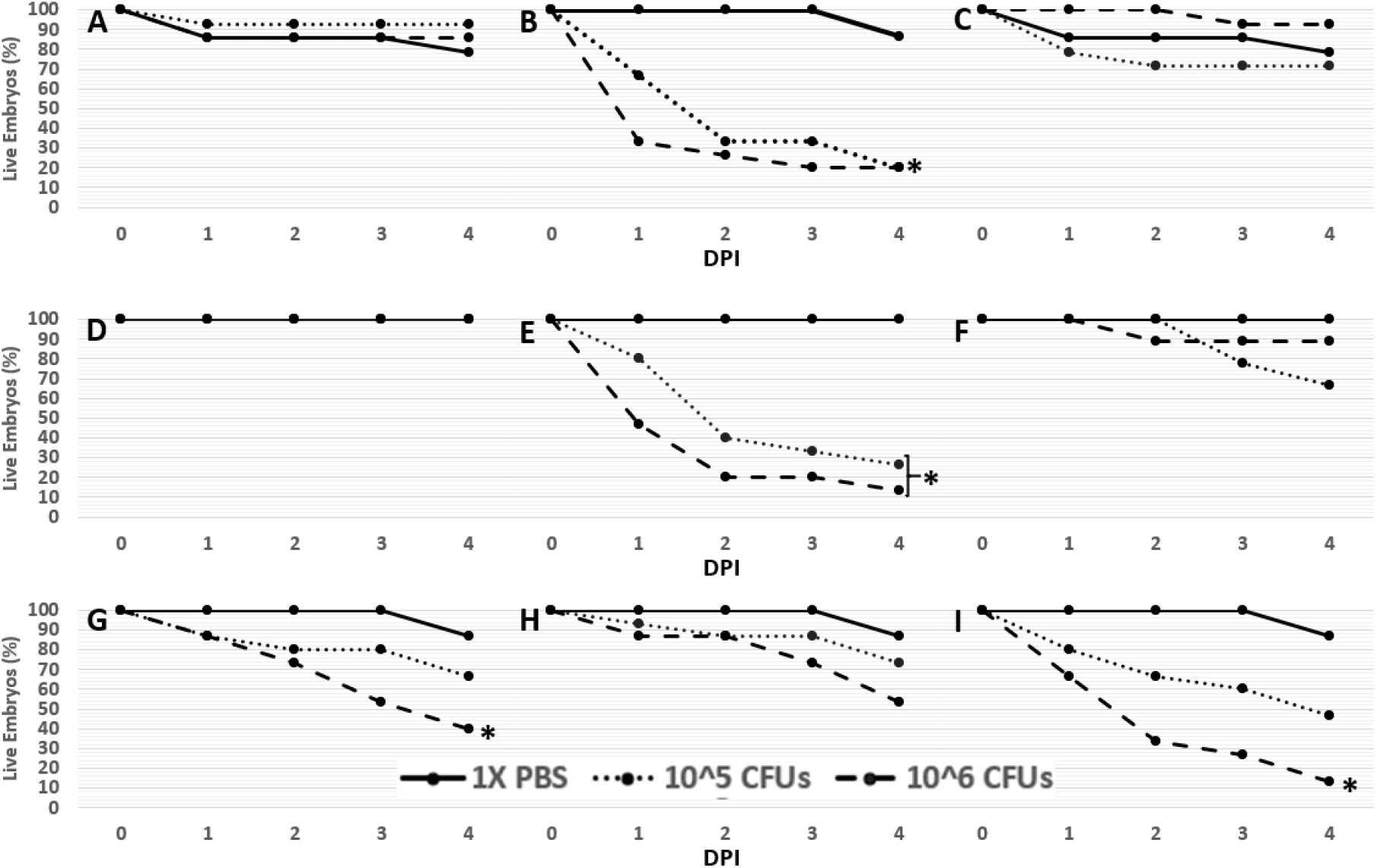
Layer chicken line embryo lethality for day post injection (DPI) with 1xPBS, 10^5^, or 10^6^ CFU of different bacterial isolates. Isolates were: A: 908 (n=14), B: 1302 (n=15), C: 1401 (n=14), D: 1409 (n=15), E: 1413 (n=15), F: 1415 (n=9), G: 1510 (n =15), H:1512 (n=15), and I: 1514 (n=15). Details are as in Figure 1. Asterisk (*) indicates treatments significantly different from 1xPBS (P <0.05) on day 4.

**Table 1:** Details of bacterial isolates utilized for ELA, including species, strain designation, host, and primary citation. In Isolate Source Lame indicates a bird number from BCO sampling. Abbreviations include: LT: Left Tibia, RT: Right tibia, LF: Left femur, and RF: Right femur.

| Species | Designation | Host | Isolate Source | Citation |
| --- | --- | --- | --- | --- |
| *S. agnetis* | 908 | Broiler | Femoral BCO; UA Research Farm | (1) |
| *S. chromogenes* | 1401 | Broiler | Thoracic Vertebrae; Lane3 | (36) |
| *E. cecorum* | 1415 | Broiler | LT/RT; Lane5 | (36) |
| *E. coli* | 1409 | Broiler | RT; Lane3 | (36) |
|  | 1413 | Broiler | Blood; Lane12 | (36) |
|  | 1512 | Broiler | LF; Lane18 | (36) |
|  | 1527 | Broiler | RF; Lane18 | (36) |
| *S. aureus* | 1510 | Broiler | LT Lane14 | (36) |
|  | 1514 | Broiler | RF Lane15 | (36) |
|  | 1302 | Human | Wound | ATCC 29213 |

We then extend the analyses to ELA using Broiler Chicken Line (BCL) embryos. As shown in Figure 3, significant embryo lethality was obtained with *S. aureus* 1302 and 1514, and *E. coli* 1413 and 1512. *S. agnetis* 908, *S. chromogenes* 1401, *E. coli* 1409, and *E. cecorum* 1415, showed no virulence for either 10^5^ or 10^6^ CFU. We did note that for all four isolates that showed lethality for BCL, both the 10^5^ and 10^6^ CFU injections showed significant embryo mortality (Figure 3 panels B, E, G, and H). For LCL, the 10^5^ injections were only different from the 1xPBS control for *S. aureus* 1302 and *E. coli* 1413. However, for the other two isolates, we might reach significance for the 10^5^ CFU injections with more embryos. We also note that lethality was more rapid in the BCL than with LCL embryos (Figures 2 and 3).

**Figure 3.**
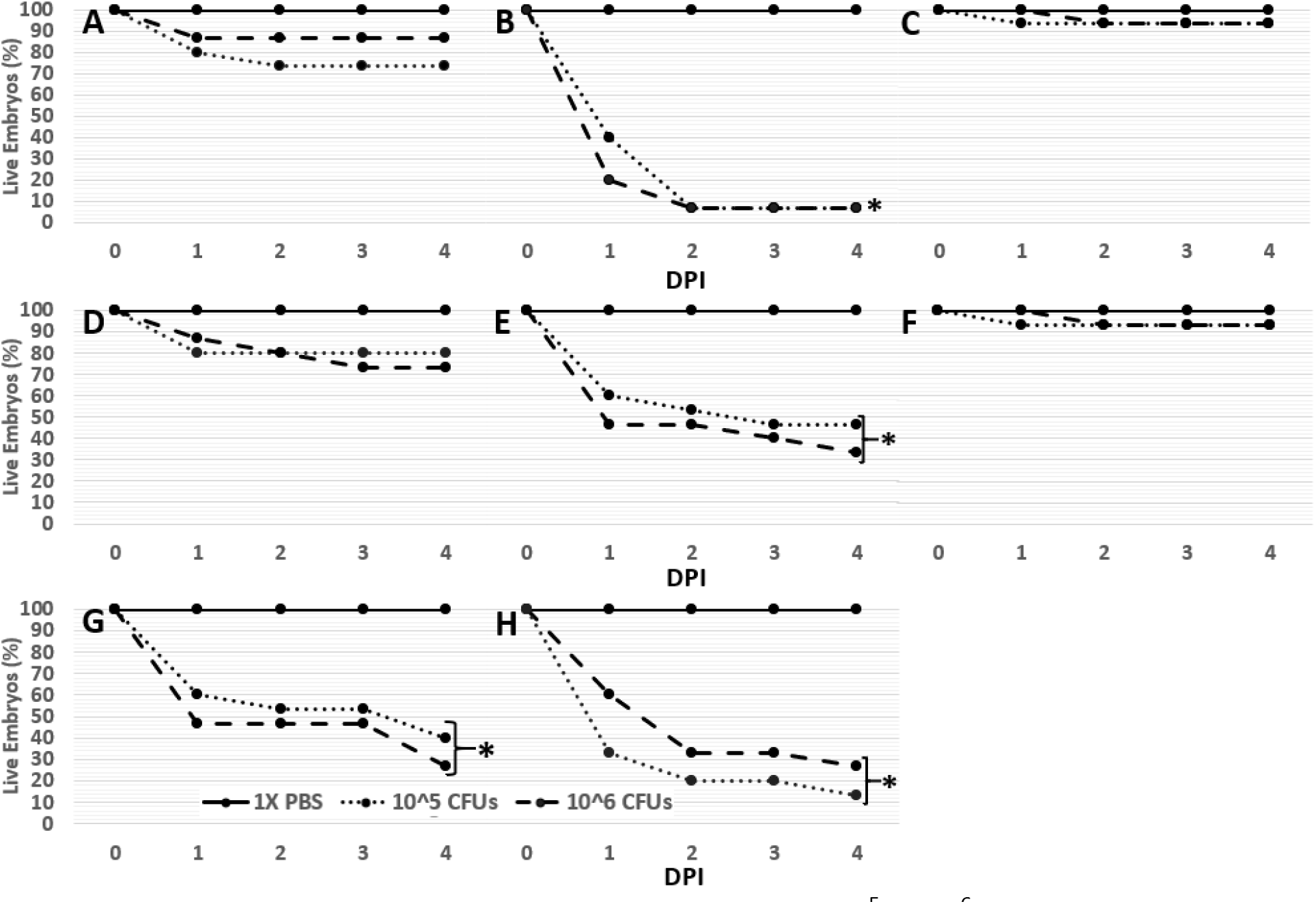
Broiler chicken line embryo lethality for injections of 10^5^ or 10^6^ CFU of bacterial isolates. Isolates were: A. 908, B. 1302, C. 1401, D. 1409, E. 1413, F. 1415, G. 1514, and H.1527. Details are as in Figures 1 and 2. For each treatment n=15.

### Repeatability of ELA as a Measure of Virulence

We noted that there was occasional variation in the ELA results for some isolates including *S. aureus* 1510 and 1514, which should be nearly identical, and *E. coli* 1409. We therefore compiled results from 11 experiments using LCL or BCL injected with 1xPBS or *E. coli* 1409 spanning nearly two years (Table 2). Embryo survival at day 5 for 1409 ranged from 100 to 11% in LCL and 73 to 44% in BCL. Close inspection of the data across experiments shows that embryo survival for 1x PBS was also varied with different batches of embryos. For the experiment of 2/20/020 the 1xPBS control was actually lower than that for 1409. We did not see this level of variability for *E. coli* 1413 as it was always highly virulent. We suspect that the variable ELA results for some isolates is likely due to differences in the particular set of eggs available for experiments. Factors affecting embryo quality might include age of hens, environmental factors in the housing (feed, heat, cold, air-quality, lighting, etc.), or post-lay conditions for specific groups of eggs (days post lay, fresh collection, etc.).

**Table 2.**
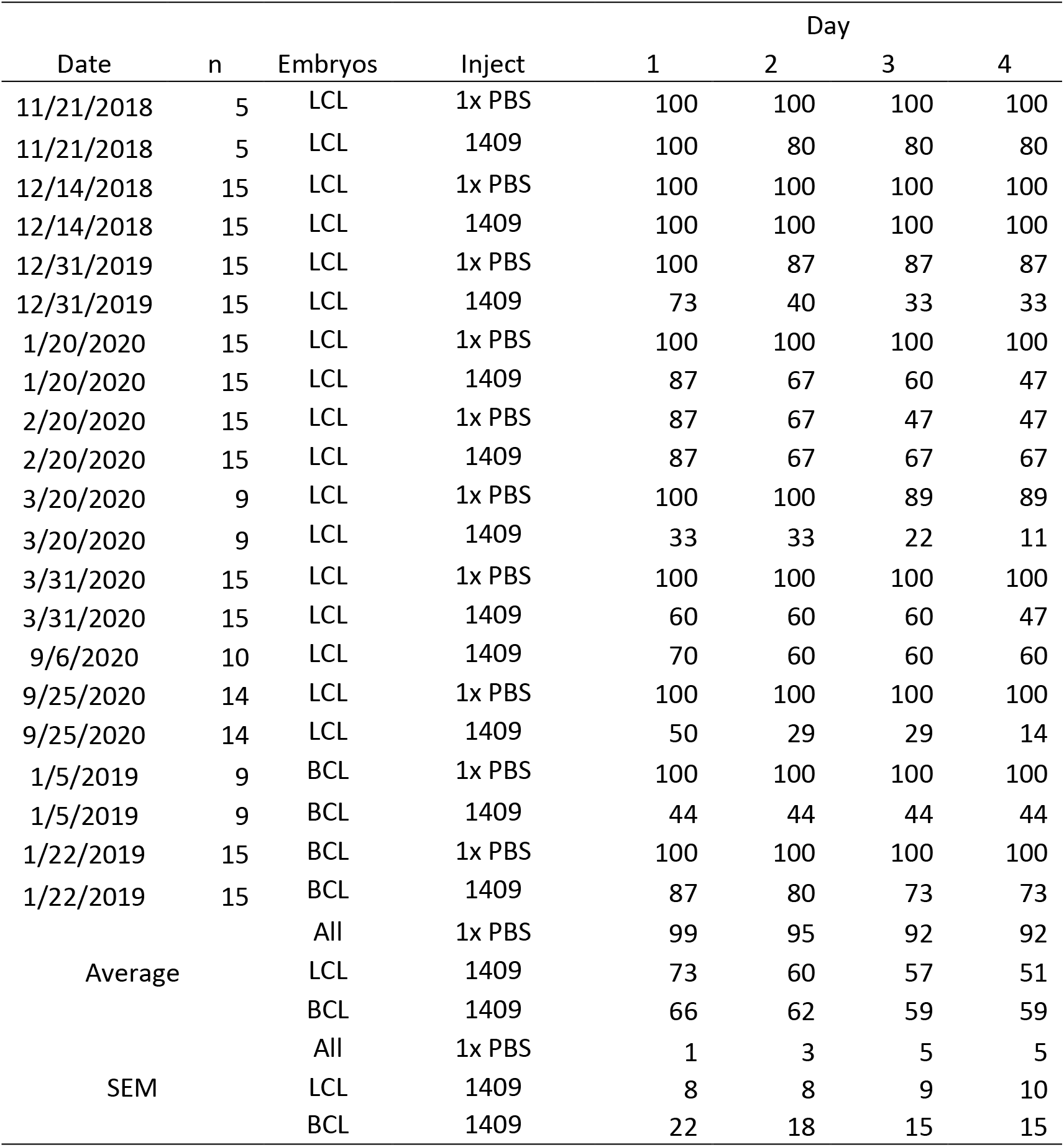
Variability of ELA results for *E. coli* 1409. On the given dates for the indicated count (n) of embryos from layers (LCL) or broilers (BCL), were injected with 100 μl of 1x PBS or 10^6^ CFU of *E. coli* 1409. Percent embryo viability is presented on the given days post injection. Averages and SEM are provided at the bottom across all experiments.

## Discussion

We performed ELA with different bacterial isolates isolated from lame broilers to estimate relative pathogenicity. We observed that *S. agnetis* 908 is not pathogenic in the ELA even though we have shown that this isolate readily infects young broilers when administered at 10^4^ or 10^5^ CFU/ml in drinking water at 20 and 21 days of age (1, 3, 6, 7). Broilers challenged with *S. agnetis* 908 begin to develop lameness by 41 days of age and by 56 days of age 50% of the birds will be clinically lame with BCO lesions in proximal femoral and tibial heads. Many of the birds develop bacteremia with hundreds to thousands of CFU/ml of blood. Additionally, the treated birds spread the infection to birds within the same room, with 30-40% of those birds lame by 56 days of age (3, 6). ELA in broiler embryos has been used to compare virulence of *E. cecorum* from BCO birds (primarily kinky back) and feces (33). Kinky back isolates showed lethality of >70% while cecal isolates showed lethality of <40%. We used *E. cecorum* 1415 collected from an infected vertebrae in a kinky back bird (36) but it showed no significant lethality (<40%) in the ELA. We compared three *E. coli* isolates from BCO lame birds (36) and found they had very different apparent virulence in the ELA. Even though these three isolates were from three different commercial broiler farms in Arkansas that were experiencing BCO outbreaks, we have shown that all 3 are very different based on whole-genome comparisons (36). This is surprising given that 1409 and 1413 were isolated on the same day from two different farms within 5 km of each other that were operated by the same integrator and supplied from the same hatchery. *S. aureus* isolates showed virulence in the ELA, including an isolate from human infection, and isolates from a BCO outbreak on a different farm operated by a different integrator. We sampled 11 lame birds from that farm and determined that 7 of the birds were infected with *S. aureus*. Genome analyses showed that the BCO *S. aureus* isolates were highly related and very closely related to *S. aureus* isolates obtained from diseased birds or broiler meat dating from 2010 in Oklahoma all the way back to the 1970s in Europe (36, 37). The clade has been identified multiple times in Arkansas and Oklahoma for at least a decade. The clade appears to be exclusively associated with poultry, so virulence in the ELA is not surprising. The genomic comparisons of *E.coli* and *S. aureus* chicken BCO isolates led us to propose that *E. coli* association with BCO is not exclusively poultry specific and that this species appears to be more of a generalist, whereas *S. aureus* and *S. agnetis* appear to be specialists and do not readily jump back and forth, infecting different host species (5, 36).

Comparison of our results with those of Borst *et al. (33)* identified several differences. They used BCO isolates and cecal isolates of *E. cecorum* in specific pathogen free (SPF) and broiler embryos. They reported that injections of 10^2^ CFU of the BCO isolates resulted in viabilities of 33% for broiler and 42% in SPF, embryos. With higher innocula the SPF embryos appeared more susceptible than the broiler embryos. Further, broiler embryos injected with 10^2^ of cecal isolates produced higher viability, circa 60%. In our experiments significant loss of viability required 10^5^ CFU. Although there was some loss of viability at lower injection levels we did not reach statistical significance for 10^3^ or 10^4^ CFU, when we used our most virulent isolate, *E. coli* 1413. Lack of significance at the lower injection quantities reflects that we used only 5 embryos per injection level. When we used more (n=15) embryos for the screens of different isolates we still had many isolates that failed to show significance when the viabilities were above 60% even with 10^5^ or 10^6^ CFU. In our hands, with our broiler embryo sources, far higher numbers of bacteria were required to reach significant levels of lethality. This difference may be because our embryos were from broilers (and layers) raised under standard conditions, whereas Borst *et al*. used embryos from SPF stocks and a broiler “flock intensively managed and has no history of *E. cecorum*-associated disease” (33). Thus, in our breeder stocks there may be higher maternal exposure to bacterial pathogens which would result in significant deposition of maternal antibodies in the eggs, which could provide enhanced immunity to the embryos. Considering that we saw marked differences in the ELA results for *E. coli* 1409 with different sets of embryos across different flocks, the quality and sources of embryos appear to be critical. Indeed, in our experiments, broiler embryos generally were more susceptible than our layer embryos. Our data suggest caution in drawing broad conclusions when using ELA for evaluation of relative virulence of BCO isolates. Concerns include: a) the lack of significant virulence by *S. agnetis* 908 which we have shown is hypervirulent in inducing lameness and spreading through a broiler facility (3, 6, 7), b) the lack of virulence of *E. cecorum* 1415 which should have been virulent based on earlier reports using ELA (33), and c) the variability of ELA results with different sets of embryos for hypo- or moderately virulent isolates (e.g., *E. coli* 1409). There remains a need to develop new, convenient, and reliable, laboratory assays for assessing pathogenicity for chicken bacterial isolates related to BCO.

## Acknowledgements

Partial support for this work was from Zinpro LLC, Chr Hansen, and the Arkansas Biosciences Institute. The funders had not input on the design or interpretations of these experiments.

AH was supported by a scholarship from the Iraqi Ministry of Higher Education and Scientific Research. AP was supported in part by a Fulbright Fellowship.

